# Olfactory epithelium local proteomic during SARS-CoV-2 infection in hamsters

**DOI:** 10.64898/2026.11.25.769107

**Authors:** Zainab Ouaid, Juliette Jacquelin, Joelton Rocha-Gomes, Claudia Bevilacqua, Carine Machado Rodrigues, Audrey Saint-Albin, Luc Jouneau, Quentin Nevers, Nicolas Meunier

**Author notes:** Authors contributed equally (Zainab Ouaid, Juliette Jacquelin, Joelton Rocha-Gomes).

## Abstract

SARS-CoV-2 has been extensively studied since its recent emergence. It has proven to be an excellent model of respiratory virus impact on nasal epithelium for its massive impact notably on the olfactory epithelium using the hamster animal model. We and others have previously observed that following infection, the nasal epithelium is massively infiltrated by innate immune cells followed by desquamation leading to release of cellular debris in the lumen of the nasal cavity. This impact varies spatially in the nasal cavity, the anterior part being infected early mainly in the respiratory epithelium followed by the olfactory epithelium in the posterior part. In order to better understand the impact on host cells, we decided to use laser microdissection to analyse the proteomic of restricted area of infected epithelium taking into account this spatial variation. We observed that protein levels are mostly decreased following SARS-CoV-2 infection at 2 days post-infection which contrasts with many genes having a stable expression level. Using available scRNASeq library, we could associate the variation of protein levels to the various cell types of the nasal cavity epithelium and our findings are consistent with a destruction of most epithelial cells and the infiltration of innate immune cells. Overall, our study confirms that SARS-CoV-2-host interactions lead to profound alteration of genes expression and that while most studies heavily rely on mRNA levels, it may be decorrelated to protein levels. Our approach offers an alternative to study host-virus interactions using spatially restricted proteomic.

## Introduction

While human smelling capacities are now recognized in par with other mammals (1), its importance in daily life appears only when this sense is impaired. Indeed, the loss of smell impacts feeding behaviour but also disrupts our ability to detect danger such as fire, gas leak or spoiled food leading potentially to depression (2). While post-viral olfactory disorders have been described for long, their origins remain elusive (3). The COVID-19 pandemic has been characterized by an unusually high level of anosmia reaching ∼50% of prevalence for the first variants and ∼10% for Omicron variants affecting millions of people (4, 5). Such high prevalence has drawn lots of interest on the impact of SARS-CoV-2 on the nasal cavity which has been explored extensively notably using hamster as an animal model (6–8). Indeed, while hamsters develop a mild disease, all infected animals suffer from olfactory disorders as early as two days post-infection (9).

Olfaction starts in the olfactory epithelium (OE) housing olfactory sensory neurons (OSN) enwrapped by sustentacular cells (SUS). Early histological studies have shown that SUS are the main SARS-CoV-2-infected cells both in hamsters and humans (8, 10). In hamsters, the infection leads to massive destruction of the OE releasing cellular debris including OSN in the lumen of the nasal cavity. Data from hamster experiments show that the infection is followed by an early infiltration of innate immune cells (7). Resident Iba1^+^ macrophages are the main innate immune cells gathering and proliferating next to the infected OE. Neutrophils are also infiltrating the tissue and play a major role in the desquamation of the infected epithelium. These events are highly variable in time and space as the infection starts in the anterior part of the nasal cavity in the respiratory epithelium then progresses to the OE in the posterior part with strong differences between the ventral and dorsal areas (6). Most molecular studies unravelling the events behind SARS-CoV-2-induced olfactory disorders are based on bulk transcriptomic or scRNAseq (11) losing the spatial resolution of these events. Only one study presents OE spatial transcriptomic data during SARS-CoV-2 infection in humans (10). While very informative, the difficulty of obtaining human biopsies does not allow to control the events occurring locally. Such technique could be used in animal studies, but the processing of OE samples preserving the integrity of the spatial organization of the nasal cavity requires relatively long processes including decalcification which may hamper the RNA quality.

Laser microdissection coupled to LC-MS/MS offers an attractive alternative. Indeed, such technique allows to selectively dissect small tissue region, keeping the spatial resolution information, and is based on protein detection which are less sensitive to degradation related to tissue processing than RNA. It has been used to resolve region-specific proteomic signatures in complex fixed tissues such as lung and liver (12, 13), but rarely in a viral context. Indeed, laser-microdissection coupled to proteomic analyses have been used to characterize the features of pulmonary damage in lungs of COVID19-deceased patients (14), but the combination of these techniques remains unexplored for the study of viral infections in animal models and at the scale of the nasal cavity, while this tissue is key for both the pathogenesis and the transmission of respiratory pathogens.

Thus, in order to locally explore the molecular events occurring in infected OE areas, we used laser microdissection coupled to LC-MS/MS to study the evolution of protein levels in the nasal cavity of SARS-CoV-2-infected hamsters.

## Material and methods

### Animals

The study was carried out following protocols approved by ANSES/ENVA/UPEC local Ethics Committee and authorized by the French Ministry of Research under the number APAFIS #35556-2022022322445750 v7 in accordance with French and European regulations. Ten specific-pathogen-free (SPF) 8-week-old male Golden Syrian hamsters *Mesocricetus auratus* (Janvier-Labs, Le Genest-Saint-Isle, France) housed under BSL-III conditions were kept according to the standards of French law for animal experimentation. Four hamsters were used as controls. Six hamsters were infected with 5.10^3^ TCID_50_ of SARS-CoV-2 isolate FRANCE/IDF0372/2020 (D614G strain circulating in Europe in early 2020) and euthanized at 2 days post-infection (dpi). The head was separated into two hemi-heads, one half was used for histology including laser microdissection and the second for RT-qPCR analysis. We did not exclude any sample and hamsters were assigned randomly to each group.

### mRNA quantification (RT-qPCR)

Total RNA was extracted from frozen nasal turbinates using the Trizol-chloroform method as previously described (7) with some modifications. Briefly, Oligo-dT first strand mixed with random hexamers cDNA synthesis was performed from 1 μg total RNA by iScriptAdv cDNA kit (Biorad, #1725038) following the manufacturers recommendations. qPCR was carried out using 125 ng of cDNA templates added to a 15 μL reaction mixture containing 500 nM primers (sequences in **Supplementary Table 1**) and iTaq Universal SYBR Green Supermix (Biorad, # 1725124) using a thermocycler (Mastercycler ep realplex, Eppendorf). Fluorescence was monitored and measured by Realplex Eppendorf Software. A dissociation curve was plotted at the end of the 40 amplification cycles of the qPCR to check the ability of theses primers to amplify a single and specific PCR product. Quantification of initial specific RNA concentration was done using the ΔΔCt method (15). Standard controls of specificity and efficiency of the qPCR were performed. The mRNA expression of each gene was normalized with the mean expression level of dicer and β-actin genes.

### Laser microdissection

Serial sections (12 μm thick) were performed on median transversal sections, perpendicular to the hard palate, highlighting the nasal structures and epithelia. Immunostaining was performed as previously described (6) on successive slides to confirm the region of interest for laser microdissection against SARS-CoV-2 N protein (1/1000; mouse monoclonal; clone 1C7C7; Sigma-Aldrich), Iba1^+^ (1/500; rabbit monoclonal; clone EPR16588; Abcam) and Arg1^+^ cells (1/200; rabbit polyclonal; PA5-29645; Thermo Fisher).

Prior to laser microdissection, cryosections were air-dried at room temperature for 5 min. Residual OCT embedding medium was gently removed by washing the sections three times with phosphate-buffered saline (PBS) for 30 s each while keeping the slides in a horizontal position to prevent tissue detachment. Approximately 500 µL of PBS was carefully dispensed to completely cover each section, and the wash solution was removed after each step by gentle decantation. Sections were then stained with a 1% (w/v) cresyl violet solution prepared in 50% ethanol for a maximum of 15 s. Following staining, the sections were sequentially dehydrated in 75% ethanol for 1 min, 95% ethanol twice for 1 min each, and 100% ethanol twice for 1 min each. Finally, the sections were cleared in 100% xylene twice for 5 min before further processing. In order to ensure appropriate dehydration, new bottles of absolute ethanol and pure xylene were opened every day immediately before use. Stained and dehydrated tissue sections were air-dried to remove xylene residues before microdissection. To obtain a perfectly flat section and minimize problems associated with microdissection, the cartilage was removed manually, using forceps, under a binocular microscope.

Selected OE areas were isolated from the whole tissue sections using an ARCTURUS XT™ Laser Capture Microdissection System and software (Applied Biosystems®, Arcturus). Laser capture was performed either under 20× or 40× magnification and infrared settings (power, duration and intensity) were adjusted for each field of view, to maximize the size of the laser spot without contaminating the sample with undesired cells. Following capture, each LCM plastic cap (CapSure®Macro LCM Caps, Arcturus) was examined at the quality control (QC) station and if necessary, undesired areas were removed from the cap by low power UV laser. After microdissection, the caps were placed directly in 0.5 ml Eppendorf tube without buffer and placed at -80°C before proteomic analysis. **LC-MS/MS**

Protein samples were processed using the iST Kit (PreOmics, Germany) according to the manufacturer’s protocol with minor adaptations for low-input samples. Briefly, samples were resuspended in 25 µL of LYSE buffer and incubated at 95°C for 30 min with agitation in an inverted microcentrifuge tube. Following brief centrifugation, proteins were digested by adding 10 µL of DIGEST solution (pH 7) and the digestion was carried out at 37°C for 3h with slow agitation. After digestion, samples were briefly centrifuged, acidified by the addition of 100 µL of STOP solution to achieve a final pH below 2. Peptides were purified using the iST cartridge according to the manufacturer’s instructions. The combined eluates were dried in a SpeedVac concentrator prior to LC–MS/MS analysis.

MS analyses were performed using a timsTOF Pro mass spectrometer coupled with a nanoElute liquid chromatography system (Bruker Daltonik GmbH, Germany). Samples were loaded on a precolumn (Acclaim PepMap C18, 0.1 mm i.d. x 20 mm, 5 μm and 100 Å pore size, Thermo Fisher) and washed with loading buffer. Then, the precolumn was connected to the separating column (Aurora C18 column, 75 μm i.d. × 250 mm, 1.6 μm, 120 Å pore size, Ion Opticks, Australia) using a two-column separation method. Buffer A consisted of 0.1% formic acid in 2% acetonitrile and buffer B of 0.1% formic acid in 100% acetonitrile. The gradient was executed at 200 nl/min with a gradient from 2% to 5% of buffer B for 1 min, 5% to 13% of buffer B for 18 min, 13% to 19% of buffer B for 7 min, 19% to 22% of buffer B for 4 min. One run took 40 min including the generation phase (95% of buffer B during 7min). The column temperature was set to 50°C. Peptides were introduced into the mass spectrometer via a CaptiveSpray nanoelectrospray ion source (Bruker Daltonik GmbH) at an electrospray voltage of 1.6 kV, an ion source temperature of 180°C, and a dry gas of 3 L/min. Data collection was carried out using dda-PASEF, briefly, a total of 6 PASEF ramps targeted precursors with a charge between 0 and 5 and a cycle time of 1.30 s. Each PASEF ramp was set to a ramp time of 180 ms. The mobility scan range used here is 0.7 to 1.10 V·s cm^2^ (1/K0), the collisional energy increased linearly from 20.0 eV at 0.60 V·s cm^2^ (1/K0) to 59.0 eV at 1.60 V·s cm^2^ (1/K0). Both MS and MS/MS spectra were recorded within a scan range of 100−1700 m/z. TIMS accumulation time was set to 180 ms, with a target precursor intensity of 14,000 arbitrary units (au) and a minimum threshold of 1000 au. Absolute intensity thresholds were set to 10 for mass spectra peak detection and 5000 for mobilogram peak detection. All samples were treated blindly of the treatment.

A FASTA format database was used from the genome of Mesocricetus auratus (MesAur1_0_pep_all) with 25912 proteins entries with the addition of N (nucleoprotein) and S (Spike) proteins from SARS-CoV-2. We also used a database of common contaminants for quality of analysis and samples handling. Database searches were performed using the X!tandem (version 2017.2.1.4) with one possible miss cleavage. Carbamidomethylation of cysteine residues and oxidation of methionine residues were set to “static” and “possible” modifications, respectively. Precursor and fragment mass tolerance was 15 ppm. Identified proteins were filtered and grouped using i2MassChroQ software (version 1.2.1). Data filtering was achieved according to a peptide E-value <0.01, protein log (E-value) <-4 and to a minimum of two identified peptides per protein (**Supp. Table 2**).

Relative quantification of protein abundances was performed using spectral counting (SC) method, defined as the number of MS2 spectra assigned to a protein. In order to select sufficiently abundant proteins, we limited our study on protein reaching a threshold of at least 15 MS2 spectra detected among the 6 samples. We then focused on proteins with a differential ratio between control and infected animals lower than 0.5 (mainly present in CTL animal) or higher than 2 (mainly present in the infected condition). STRING-db analysis was performed to cluster these proteins by ontology (0.4 interaction score; 5% FDR stringency; Text mining and Co-occurrence protein links excluded).

### scRNAseq data set analysis

A previous scRNA-eq data set of hamster OE was analysed using Seurat package (11). **Supplementary data 1** details the workflow applied to raw data. We defined 22 clusters that we reduced to 16 by gathering 4, 3 and 2 different clusters for sustentacular cells, mature and immature olfactory neurons respectively. We then used these clusters to define cell type associated proteins from our proteomic results (**Supp. Fig. 1**).

### Statistical analysis

Mann-Whitney statistic tests were performed to assess the difference between groups using Prism 8.0 (GraphPad). In graphs, error bars indicate the SEM. Pearson correlation on proteomic quantification values between samples and plot were done using corrplot R package. Detailed information on statistical test used, sample size and *P* values are provided in the figure legends.

## Results

### Laser microdissection of infected olfactory epithelium

We performed local proteomics of SARS-CoV-2-infected epithelium using laser microdissection coupled with LC-MS/MS (**Fig. 1**). We focused on the posterior part of the nasal cavity containing mainly OE (**Fig. 2A**). We used the histological changes of the OE to select infected areas. This epithelium is heavily infected at 2 dpi with typical local innate immune cell infiltration (**Fig. 2D**) leading to epithelium desquamation and cellular debris filling the lumen of the nasal cavity. We microdissected three similar regions of OE from infected and non-infected animals (**Supp. Video 1**). To assess the reproducibility of the results, the 3 samples were taken from two different animals for each condition. The two slices originating from the same animal (“2.1” and “2.2”) were separated by 120 µm. For each sample, we collected around 1/4 of the OE available in a coronal section corresponding to ∼2 mm^2^ of OE along cellular debris in the lumen of the nasal cavity. The presence of the SARS-CoV-2 N protein as well as infiltration of Iba1^+^ macrophages were confirmed by immunohistochemistry on successive slices (**Fig. 2B-C-D**).

**Figure 1:**
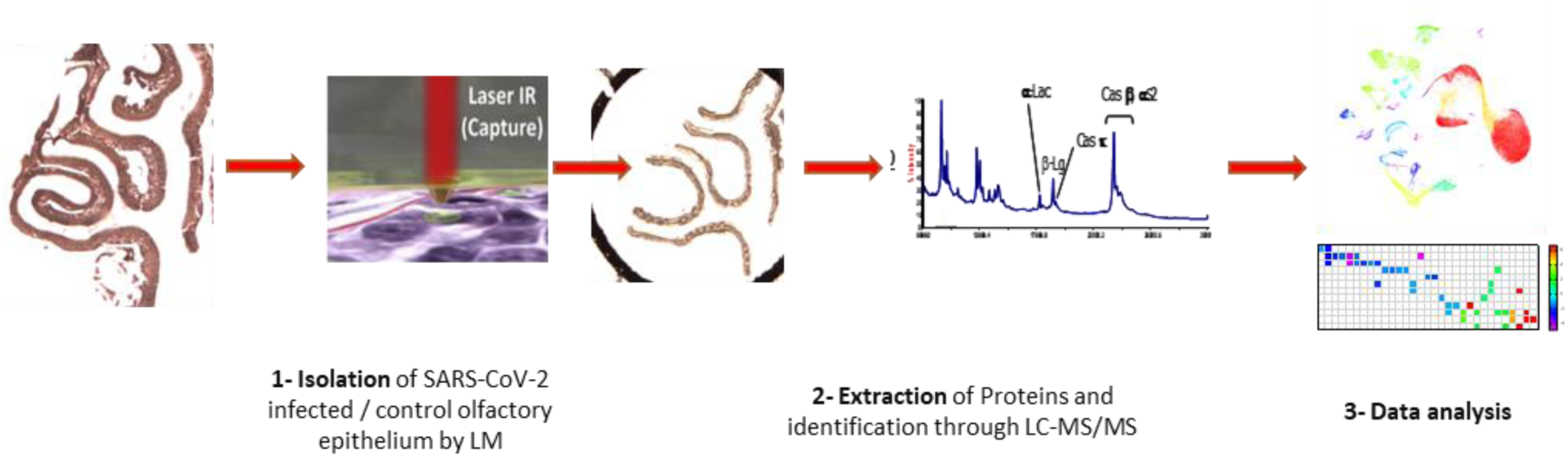
Workflow of laser microdissection (LM) of the nasal epithelium coupled to LC-MS/MS. In this study, we laser microdissected slices from the nasal cavity of 2 days post-SARS-CoV-2 infected or control hamsters in order to isolate the infected olfactory epithelium and similar regions in control animals (n=3 for each condition from 2 animals for each condition). The samples were then analysed by LC-MS/MS.

**Figure 2:**
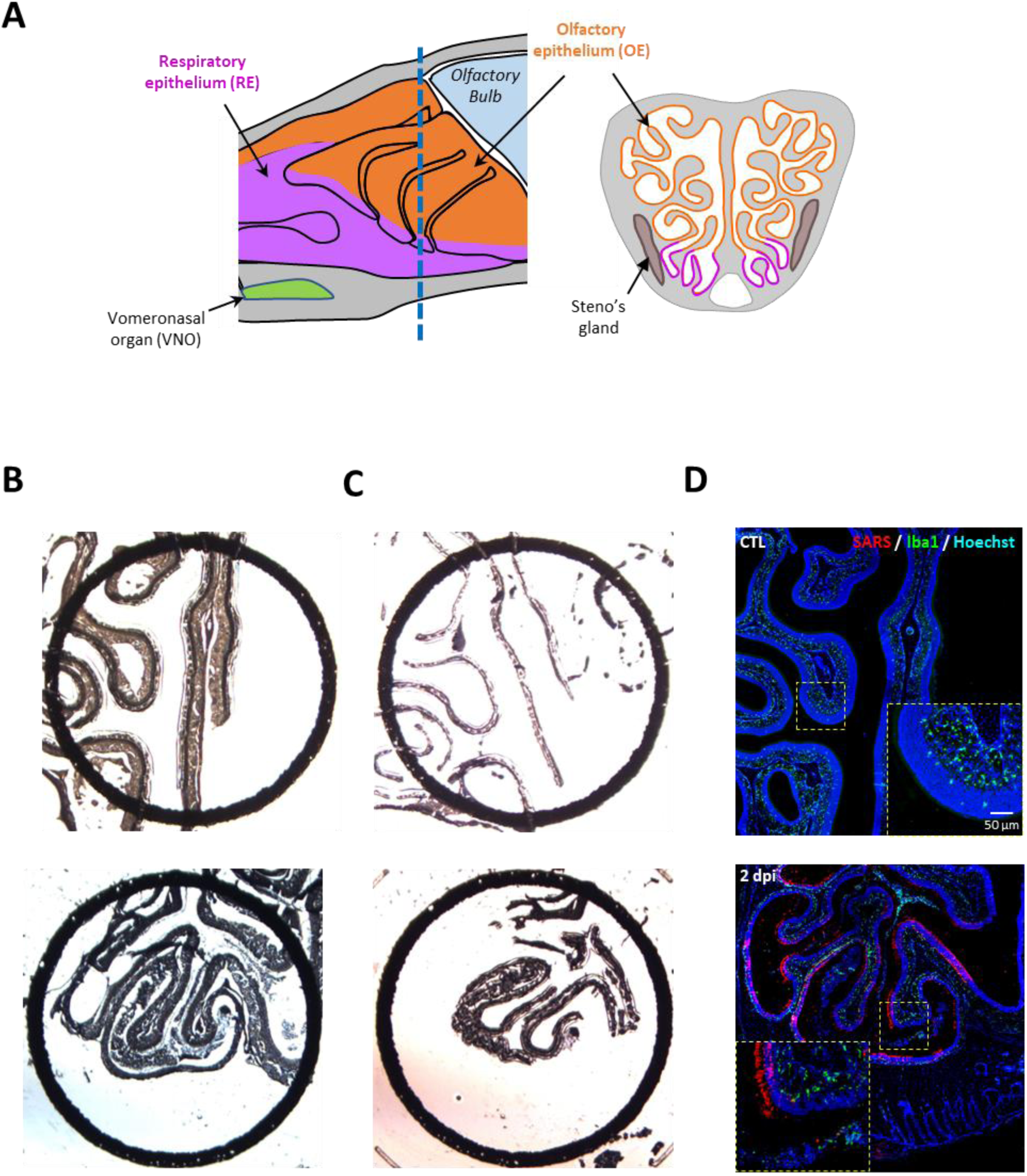
Microdissected tissue localization. (**A**) Schematic view of the hamster nasal cavity showing the coronal section used in this study. (**B**) Representative images of the microdissected area before or (**C**) after laser microdissection (LMD). (**D**) Immunostaining performed on a successive slice used for LMD. Up: non-infected animal -CTL-, Down: 2 days post-infection animal -2 dpi-showing presence of the SARS-CoV-2 N protein (red), Iba1^+^ resident macrophages (green) in the same area displayed on **B** & **C**.

### Change of local protein levels in infected olfactory epithelium

LC-MS/MS was performed on each sample separately using *Mesocricetus auratus* database as well as SARS-CoV-2 N and S protein sequences. We detected 1747 different proteins across all samples (**Supp. Data 1**). We focused our analysis on ∼1/3 of most abundant protein corresponding to 698 proteins above a threshold of at least 15 occurrences among all 6 samples. For each protein, we calculated the ratio of peptide fragment detected between infected and control samples. 456 proteins were detected with a ratio of peptide fragment occurrence between 0.5 and 2. We focused our analysis on proteins with a ratio inferior to 0.5 (208 including Pck2 and Hpgd only present in non-infected samples); an infected/non-infected ratio superior to 2 that were either present in both conditions (26) or only in infected samples (9 including N and S Spike proteins) (**Fig. 3A; Supp. Fig. 2**). These 3 categories defined protein levels respectively at least “50% lower”, “100% higher” in infected samples compared to non-infected one or “only present” during infection.

**Figure 3:**
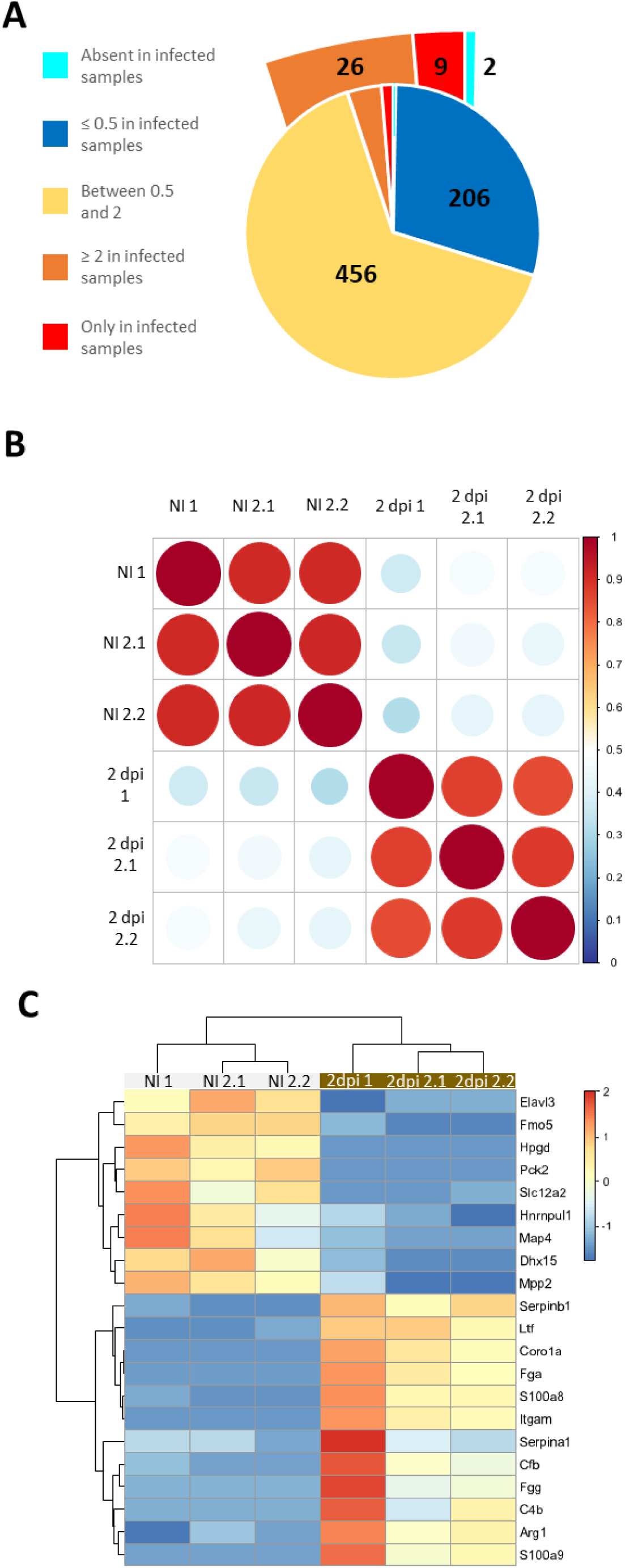
Properties of proteins detected after LM-LC/MS-MS. (**A**) Repartition of proteins according to their presence or absence in infected samples as well as their ratio between infected at 2 days post infection (dpi) and CTL samples (NI for non-infected). (**B**) Pearson correlation plot of protein abundancy between samples. (**C**) Z-score based heatmap of a selection of proteins detected across the samples.

We first globally compared the correlation of protein levels between samples without a priori (**Fig. 3B**). Pearson correlation plot shows that among all 6 samples, the CTL and 2 dpi conditions were clearly segregated (less than 50% correlation between condition compared to more than 85% within each condition). For each condition, the samples originating from the same animal were highly correlated (91% and 88% for non-infected and infected animals respectively). We next normalized by a z-score the variability of abundance of the different proteins allowing to construct an heatmap based on hierarchical clustering of the proteins among samples. This heatmap shows a similar evolution of the different protein levels between samples originating from the same animal and of the same condition (**Fig. 3C and Supp. Fig 3**). Such representation clearly shows that the OE infection by SARS-CoV-2 leads mainly to a local decrease of host protein levels consistent between animals and samples from the same animal.

### Proteomic *versus* transcriptomic levels

We next compared the level of proteins with the related genes expression measured by RT-qPCR. We selected all genes related to the proteins of the “only in infected samples” group as well as a similar number of genes related to proteins with the highest differential ratio between CTL and 2 dpi in each category (**Fig. 4**).

**Figure 4:**
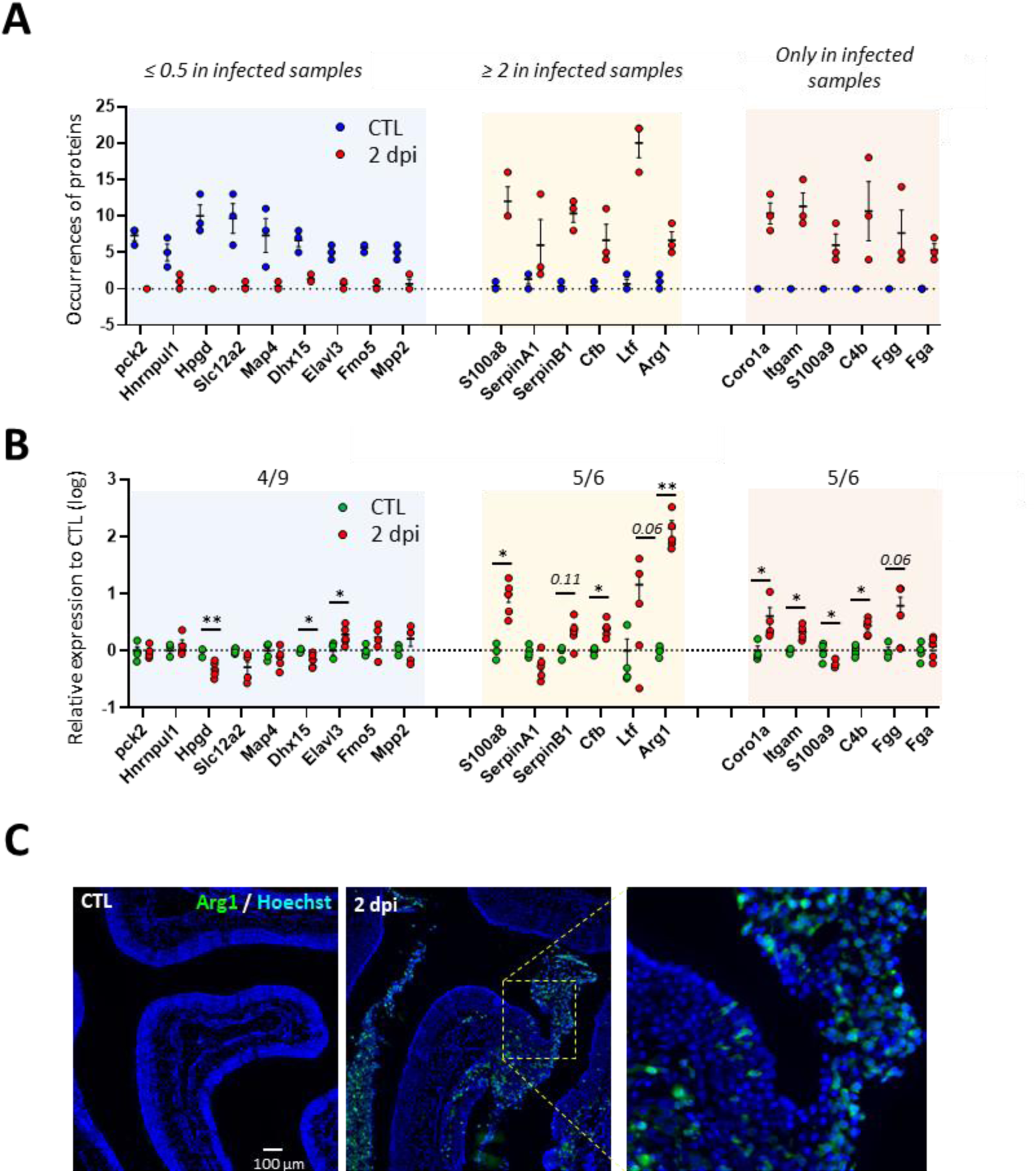
Comparative evolution of protein levels detected after laser microdissection *versus*genes expression level in olfactory turbinates. (**A**) Levels of protein detected in each sample (Mean ± SEM; n=3). (**B**) Relative expression of the corresponding genes in olfactory turbinates of non-infected (CTL) or 2 days post-infected (2 dpi) animals. For each protein category, the number of genes with a differential level of expression between CTL and 2 dpi animals following a similar difference (higher or lower) is indicated. (Mean ± SEM; n≥4; Mann-Whitney test; *P < 0.05; **P < 0.01). (**C**) Representative images of Arg1 immunostaining in similar coronal section used for LMD. Nasal cavity from non-infected animal -CTL-and 2 days post-infection animal -2 dpi-.

We detected genes expression for all genes except for MPO which is known to be only expressed in neutrophils during haematopoiesis (16). Globally, we observed strong discrepancies when comparing mRNA and protein levels in the group of proteins with a ratio ≤ 0.5 in infected samples. In this group, the related gene expression levels did not change much and only 4 out of 9 had a tendency to diminish (with only 2 reaching a statistically significant decrease). On the contrary, protein and related gene expression levels evolution were mostly consistent for the group with a ratio “≥ 2 in infected samples” and the group “only present in infected samples”. For both, 5 out of 6 gene expression levels got a tendency to increase (all of them reaching or nearly reaching statistically significance). We confirmed by immunostaining the appearance of Arg1^+^ cells in the infected area of the OE (**Fig. 4C**). Using STRING-db (17), we examined the ontology of the differentially present proteins (**Fig. 5**). Among the 241 proteins analysed, we found 13 clusters gathering 222 proteins without a clear separation by the ratio of presence between the 2 dpi and CTL group with the exception of the cluster “complement and coagulation cascades” with a ratio of 39 (**Supp. Table 4**).

**Figure 5:**
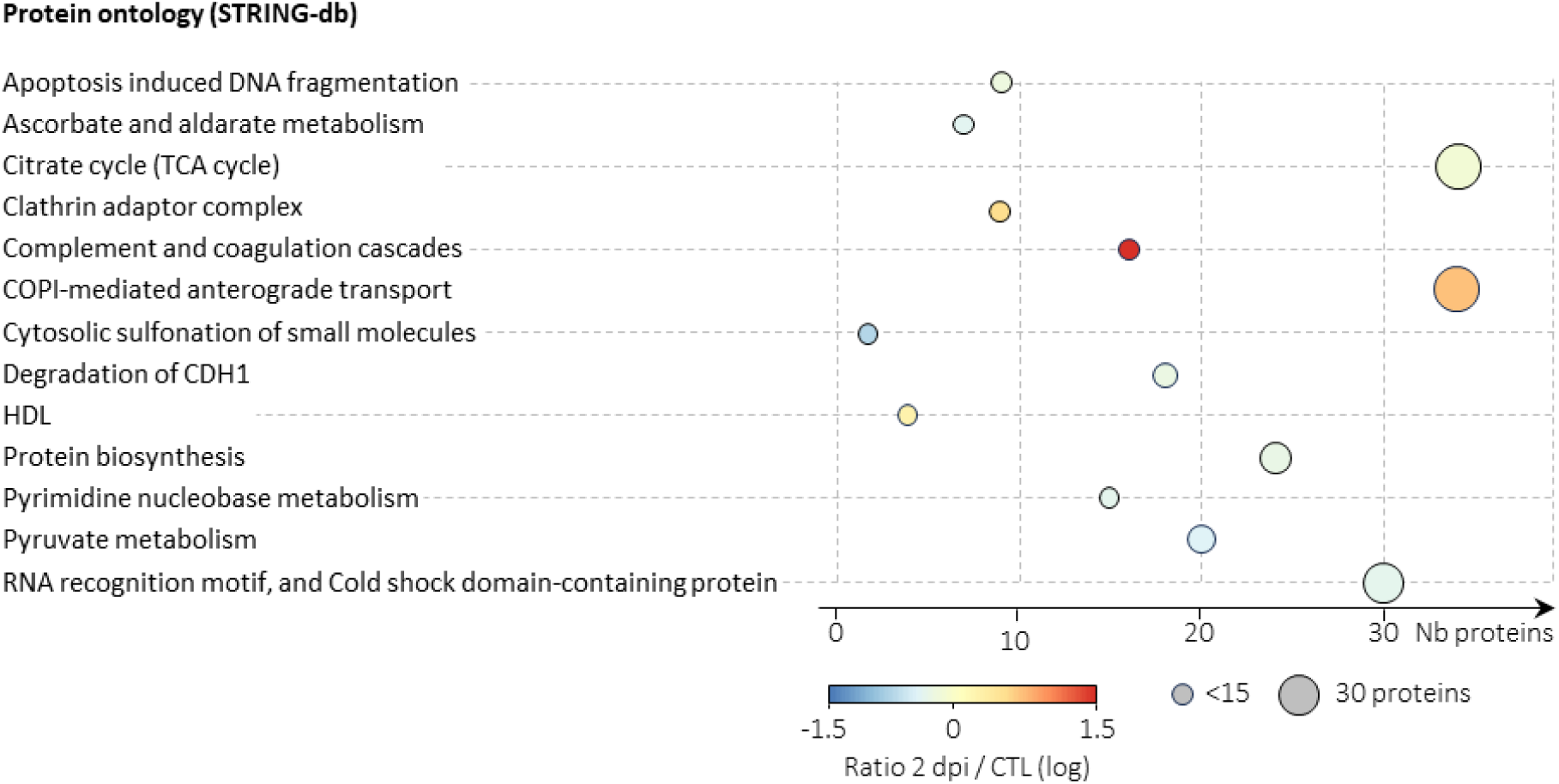
Ontology of differentially present proteins between infected and CTL sample. STRING-db analysis showing the different ontology clusters.

### Modulation of protein levels and cellular identity

We next explored whether some of the protein changes could be associated to a given cell type in the OE. We used an available scRNAs eq dataset from control and SARS-CoV-2 infected OE in hamsters (11) allowing us to define different cell clusters corresponding to our study conditions. We obtained 14 identified cell types and two unspecified clusters (**Fig. 6A**). For each cell type, we explored the level of expression of genes corresponding to proteins differentially present in the 3 groups (243 proteins, **Supp. Data 2**). We then selected the top 6 expressed genes and kept only the genes expressed in less than 4 different cell types. Such approach defined “associated proteins” to a given cell type. We then constructed a table with the different cell types and associated protein levels (**Fig. 6B**). We included MPO present in the “only in the infection samples” group as this protein is known to be specific to neutrophils but MPO gene expression is not detected in mature cells. Overall, with such filters, we associated 30 proteins differentially present between CTL and 2 dpi to 12 cell types. Seventeen of them were only found in a given cell type such as Atp2a3 in microvillar cells and 12 in only two different cell types. These cells could be classified in 3 groups (i) associated mostly with protein level decreasing in infected samples (Microvillar cells, olfactory neurons (immature and mature), sustentacular cells, HBC) (ii) associated to protein with an increasing ratio in infected samples (macrophages / monocytes; neutrophils and lymphocytes) (iii) a mix (GBCs, NK cells, OECs).

**Figure 6:**
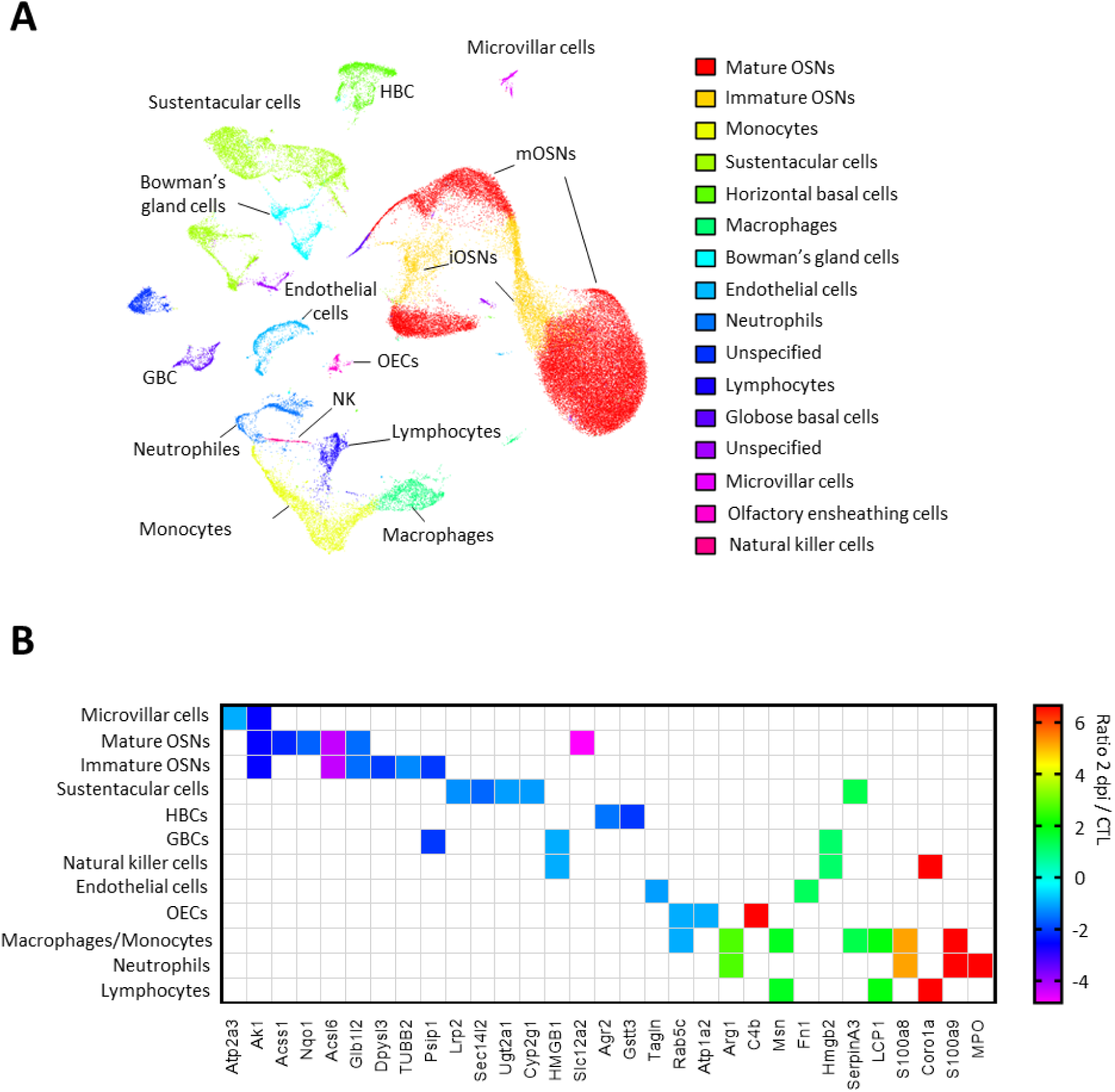
Proteins associated to specific cell populations. (**A**) UMAP plots of hamster olfactory epithelium for mock and 3 days post infection (dpi) animals (HBC/GBC: horizontal / globose basal cells; mOSNs / iOSNs: mature / immature olfactory sensory neurons; OECs: olfactory ensheathing cells; NK: Natural killer cells. (**B**) Cell type associated protein level ratio between 2 days post infection and non-infected animals

## Discussion

While the nasal cavity represents a critical entry portal and replication site in the pathogenesis and transmission of clinically-important respiratory pathogens (including human coronaviruses, influenza viruses, and respiratory syncytial virus (RSV)), the virus-induced host response within this specific tissue remains incompletely understood (18). Standard bulk transcriptomic studies and single-cell RNA sequencing (scRNAseq) approaches have yielded invaluable insights into viral pathogenesis; however, they inherently lack spatial context and/or require tissue dissociation steps that can alter cell states (19). Although spatial transcriptomics has recently emerged to bridge this gap, long tissue fixation especially required when working with BSL3 pathogens can compromise mRNA integrity. This issue is aggravated when working with tissue including bones such as the nasal cavity that requires long decalcification processes. Furthermore, transcripts abundance often correlates poorly with functional protein levels due to complex post-transcriptional and post-translational regulatory mechanisms (20, 21). To overcome these limitations, we implemented a spatially resolved proteomic workflow combining laser microdissection and high-sensitivity mass spectrometry to quantify protein level changes in infected areas of the OE. We worked with an ancestral SARS-CoV-2 strain (a variant that we had previously well characterized for its pathophysiology in the nasal cavity) and observed the presence of N and S SARS-CoV-2 proteins only in infected samples with a superior presence of the N protein (**Supp. Fig. 2**) as expected. The results were also consistent among the samples showing the robustness of our workflow (**Fig. 3B**). The presence of some of these proteins are limited to the infected samples, such as MPO and Arg1 related to neutrophils (16) and M2 macrophages respectively (22). We already published by immunostaining that MPO signal only appears in infected epithelium area (7) and we confirmed here that it holds true as well for Arg1 (**Fig. 4C**) supporting our approach.

While most host-response studies primarily focus on innate immune activation and up-regulated pathways such as interferon signalling, our proteomic analysis revealed a predominant down-regulation of protein abundance within the OE following SARS-CoV-2 infection (**Fig. 4**). Interestingly, this pronounced depletion at the protein level was rarely mirrored at the transcripts level in our targeted RT-qPCR experiments. This divergence strongly suggests the involvement of post-transcriptional silencing, accelerated protein degradation, or viral host shutoff mechanisms (21), underscoring the necessity of direct proteomic quantification as a complementary approach to RNA-based technologies. The protein depletion is also consistent with the destruction of the OE following SARS-CoV-2 infection as the diminishing protein could be linked to the loss of OSNs, SUS, microvillar and basal cells representing most of OE cells (**Fig. 6B**). Using STRING-db protein ontology, we saw that one cluster was mainly present in the infected samples (**Fig. 5**). This cluster corresponds to “complement and coagulation cascades” consistent with the increased presence of blood-related proteins which can be linked both to increased endothelial permeability during diapedesis as well as damage of the tissue following infection. Among the clusters, the “degradation of CDH1” is linked with epithelium remodelling (24) consistent with the damage of the infected epithelium.

Among the proteins displaying the most striking suppression, phosphoenolpyruvate carboxykinase 2 (PCK2) and 15-hydroxyprostaglandin dehydrogenase (HPGD) became entirely undetectable in SARS-CoV-2-infected samples. Interestingly, this loss correlated with a marked decrease in mRNA levels for HPGD, whereas PCK2 transcript levels remained largely unaltered, highlighting distinct levels of regulatory control. PCK2 is a pivotal mitochondrial enzyme involved in various metabolic pathways, whose significance has been mostly documented in cancer biology but not in the context of viral infections (25). Although its role in SARS-CoV-2 infection is not characterized, the complete loss of PCK2 in infected tissues may reflect mitochondrial dysfunction, elevated mitophagic turnover, or a viral-induced reprogramming of host metabolic flux. Conversely, HPGD serves as a key enzyme in the catabolism and inactivation of pro-inflammatory prostaglandins, notably prostaglandin E2 (26). In the respiratory tract in particular, HPGD acts as an important metabolic checkpoint to restrain vasodilation and excessive immune recruitment. The strong down-regulation of HPGD observed in our model may be linked to local hyper-inflammation, tissue oedema or other acute lesions characteristic of SARS-CoV-2 infection. However, as for PCK2, links between HPGD and viral infections are currently lacking. Thus, further *in vitro* mechanistic studies using cultured epithelial cells or organoids will be required to decipher how SARS-CoV-2 targets these proteins. In contrast to these depleted markers, we identified a subset of proteins that were strongly up-regulated or even exclusively detected upon infection. For these proteins, the correlation with RT-qPCR mRNA levels was substantially higher. Functional annotation indicated that these enriched markers primarily correspond to infiltrating immune cell populations (such as neutrophils and monocyte-derived macrophages) responding to the local inflammation.

Overall, our study shows the interest of proteomics performed on sub areas of an infected tissue. Our LC-MS/MS spatial proteomics technique could be applied to compare host proteomic shifts elicited by ancestral versus Omicron variants in order to decipher how host-virus interactions can evolve. One limitation of our study is the resolution which is limited to sub area of the nasal cavity epithelium. We could indirectly link the variation of protein level to a limited number of cell type thanks to scRNASeq available database. While our result is very consistent with the described impact of SARS-CoV-2 infection of the OE resulting on the destruction of OE cells and massive infiltration of immune cells, it is biased as RNA and protein levels are not correlated. It could explain why some cell types such as natural killer cells are associated with proteins with opposite ratio (**Fig. 5B**). The recent advances in laser microdissection technology coupled with increased sensitivity of proteomics should rapidly allow to perform single cell proteomic in infected samples unravelling new sides of host virus interactions (27).

## Acknowledgements

We thank Sophie Le Poder and Bernard Klonjkowski (UMR 1161 Virologie, Ecole nationale vétérinaire d’Alfort) for their help with BSL3 experiment.

## Author Contributions

Conceptualization: NM; QN; CB; CMR. Investigation: ZO; JJ; JRG; ASA; CB; CMR; and NM. Writing: NM and QN, with input from all authors.

---

**Table 1:**
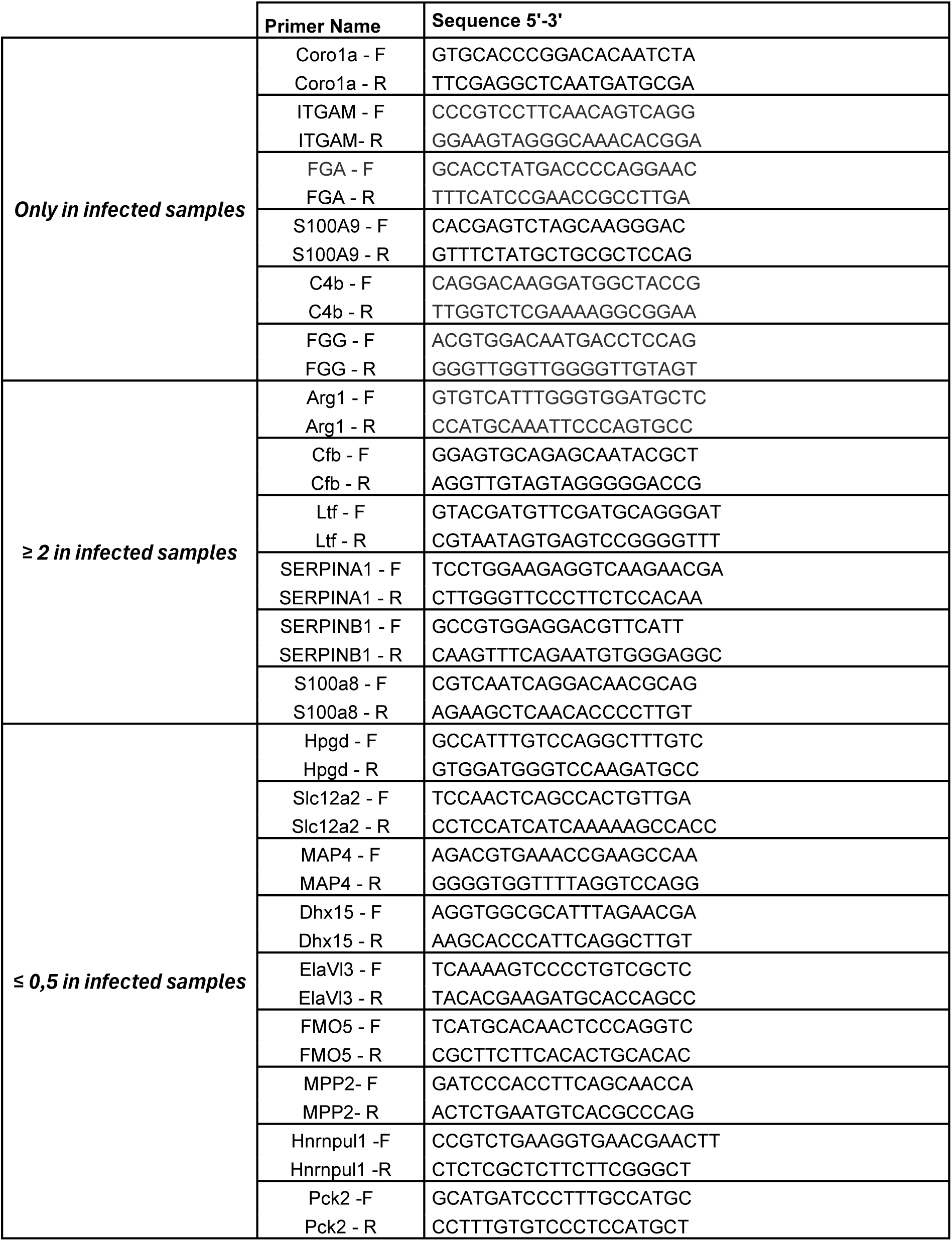
List of primers.

**Supplementary Figure 1:**
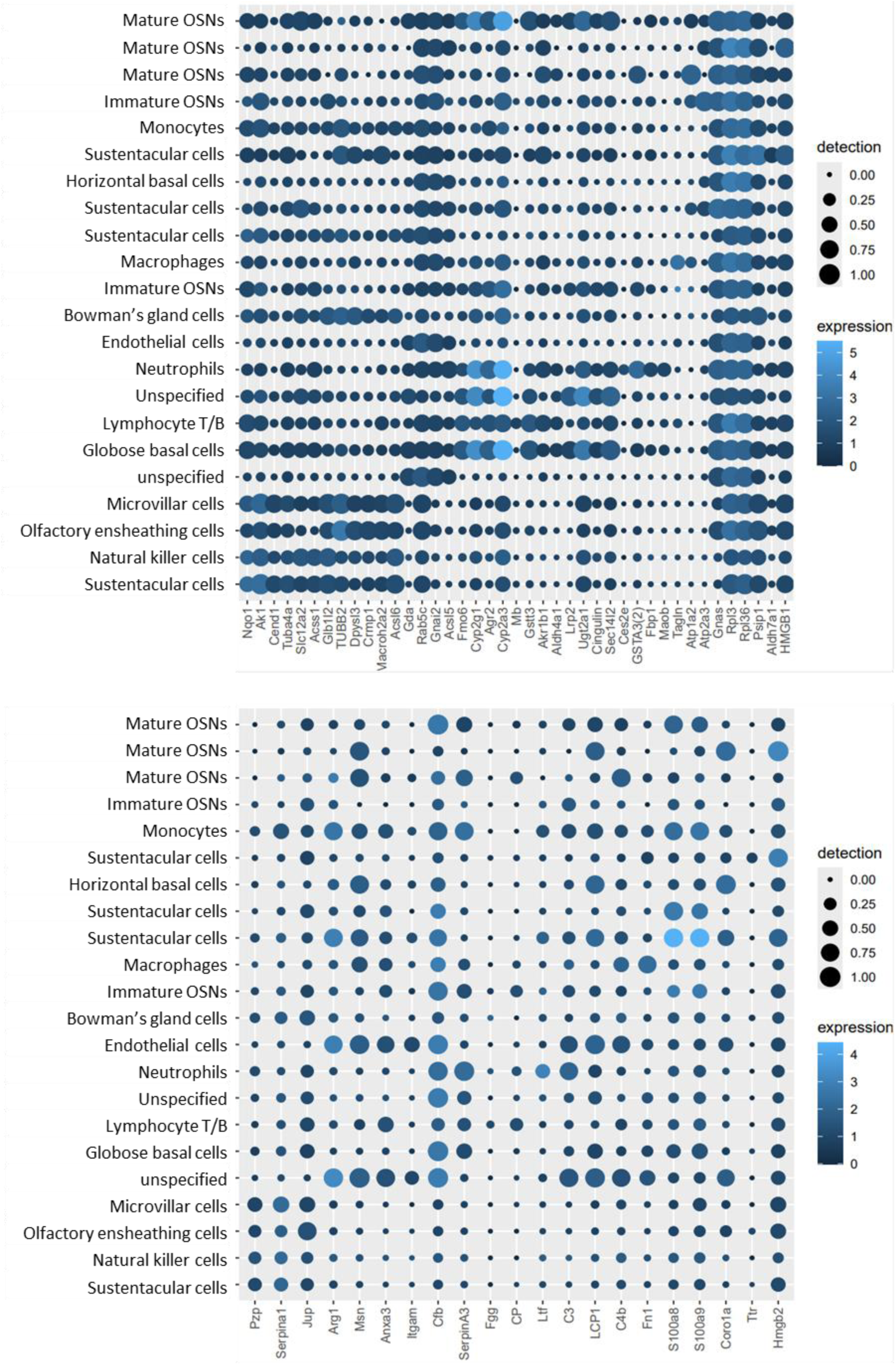
Bubble plot showing expression and presence levels in the different scRNASeq clusters of the proteins differentially expressed between control and infected samples

**Supplementary Figure 2:**
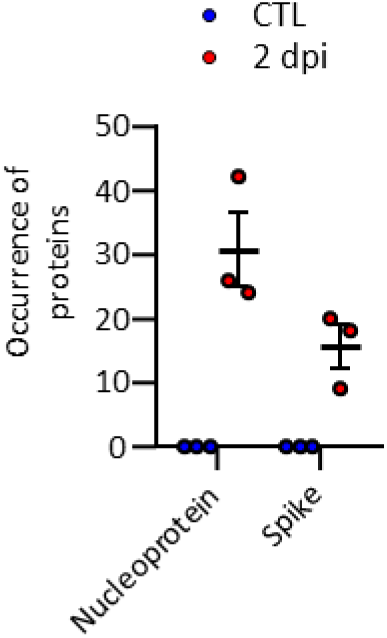
SARS-CoV-2 nucleoprotein and spike protein occurrences detected in control or infected samples (Mean +SEM; n=3)

**Supplementary Figure 3:**
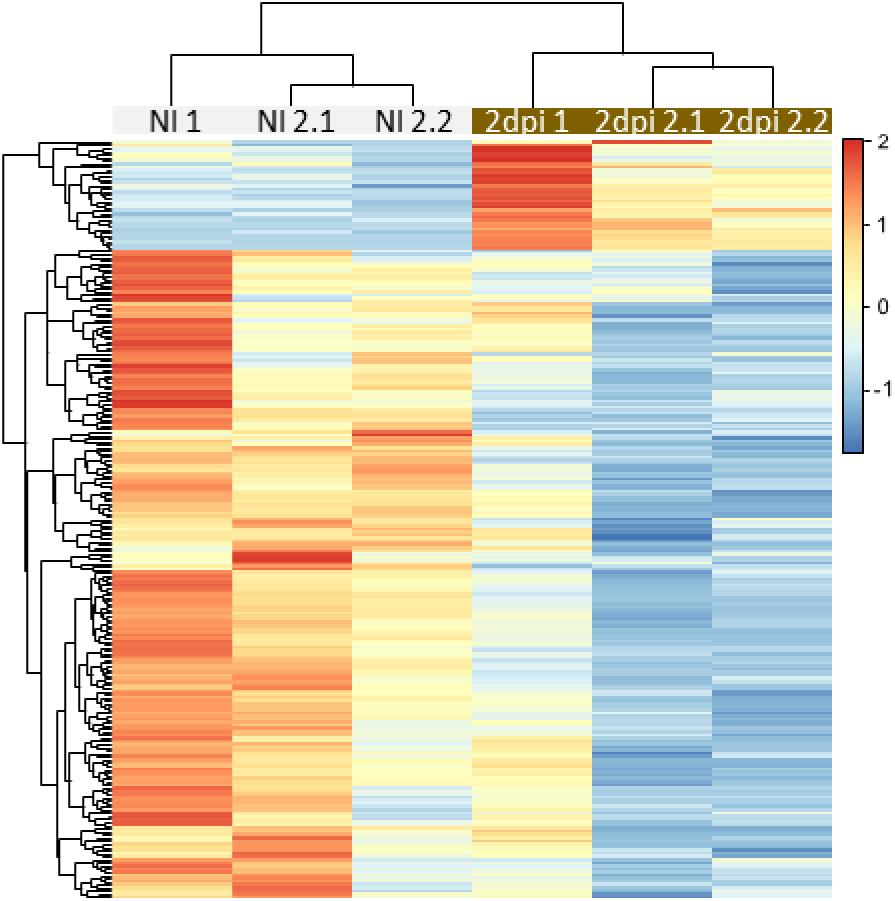
Z-score based heatmap of allproteins detected across the samples similarly as Fig. 3C

**Supplementary Figure 4:**
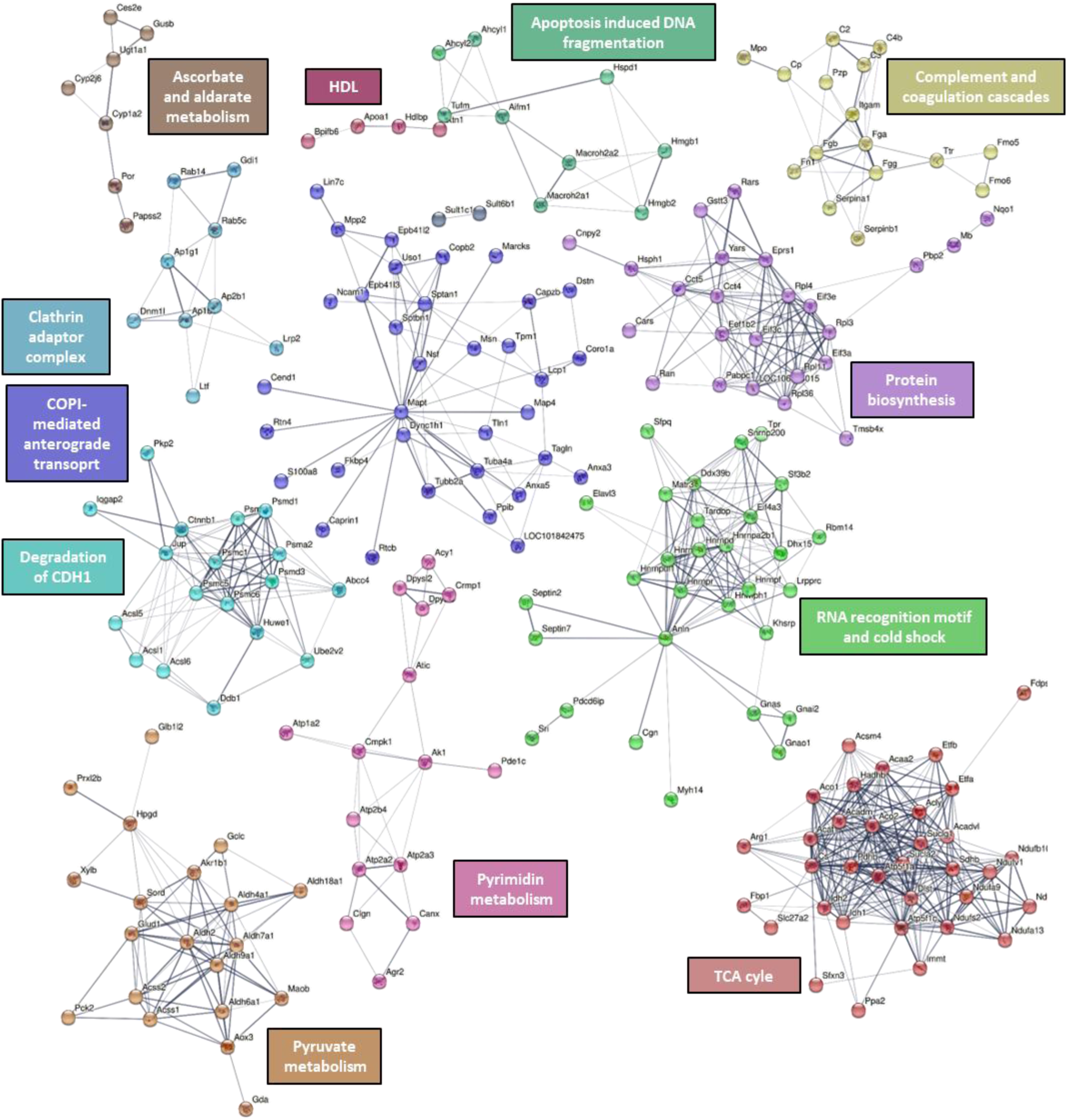
STRING-db analysis of proteins thatare differentially present following SARS-CoV-2 infection in the laser microdissected area

**Supplementary Data 1: Workflow used for analyzing single cell RNA-Seq samples**

**1°) Quality control -Filtering parameters:**

We kept cells satisfying the following quality criteria

| **Library size** | **# of genes detected / cell** | **Max % of mitochondrial expression / cell** | **Top variable genes** |
|---|---|---|---|
| [1000 ; 20000] | [1000 ; 20000] | 25 | 1000 |

**2°) Dimension reduction:**

**Gene selection**

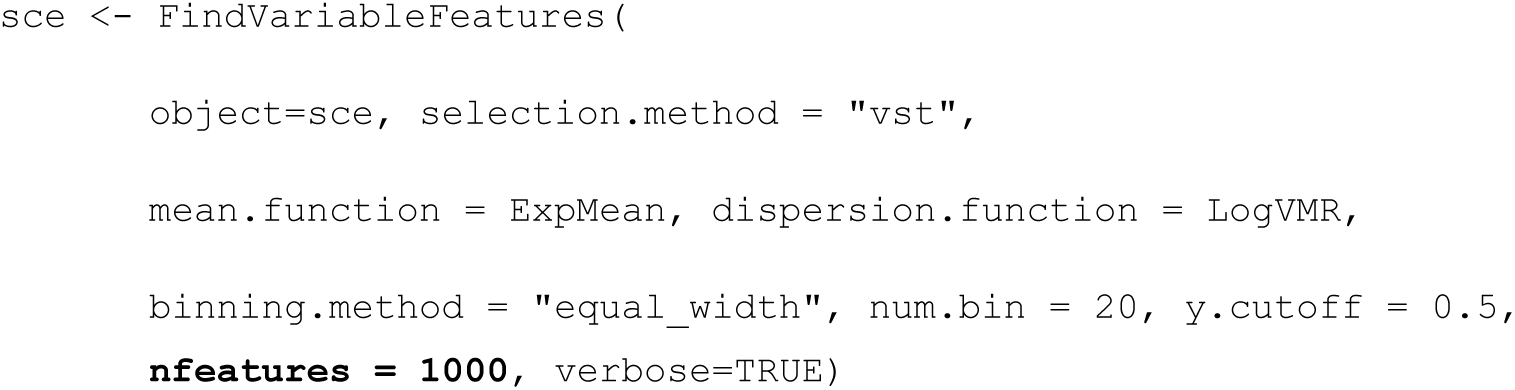

PCA components considered (runPCA Seurat function)

| Sample | PCA components considered |
|---|---|
| 0dpi_rep1 | 15 |
| 0dpi_rep2 | 15 |
| 1dpi_rep1 | 10 |
| 1dpi_rep2 | 12 |
| 3dpi_rep1 | 19 |
| 3dpi_rep1 | 19 |
| 10dpi_rep1 | 15 |
| 10dpi_rep2 | 15 |

**3°) Clustering and UMAP**

**Cluster resolution**

| **Sample** | **Cluster resolution** |
|---|---|
| 0dpi_rep1 | 0.3 |
| 0dpi_rep2 | 0.4 |
| 1dpi_rep1 | 0.4 |
| 1dpi_rep2 | 0.4 |
| 3dpi_rep1 | 0.5 |
| 3dpi_rep1 | 0.4 |
| 10dpi_rep1 | 0.3 |
| 10dpi_rep2 | 0.3 |

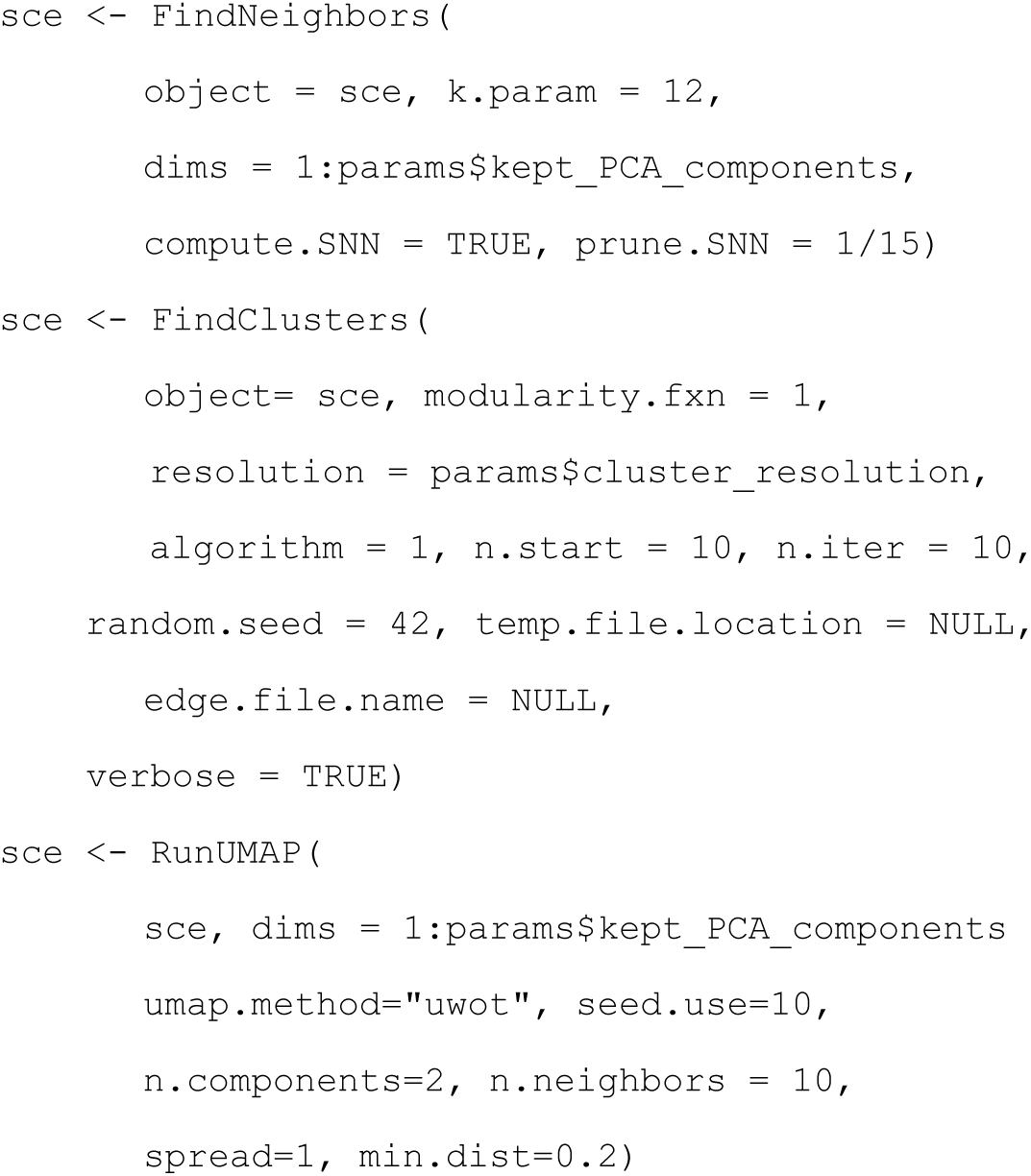

**4°) Integration of all samples:**

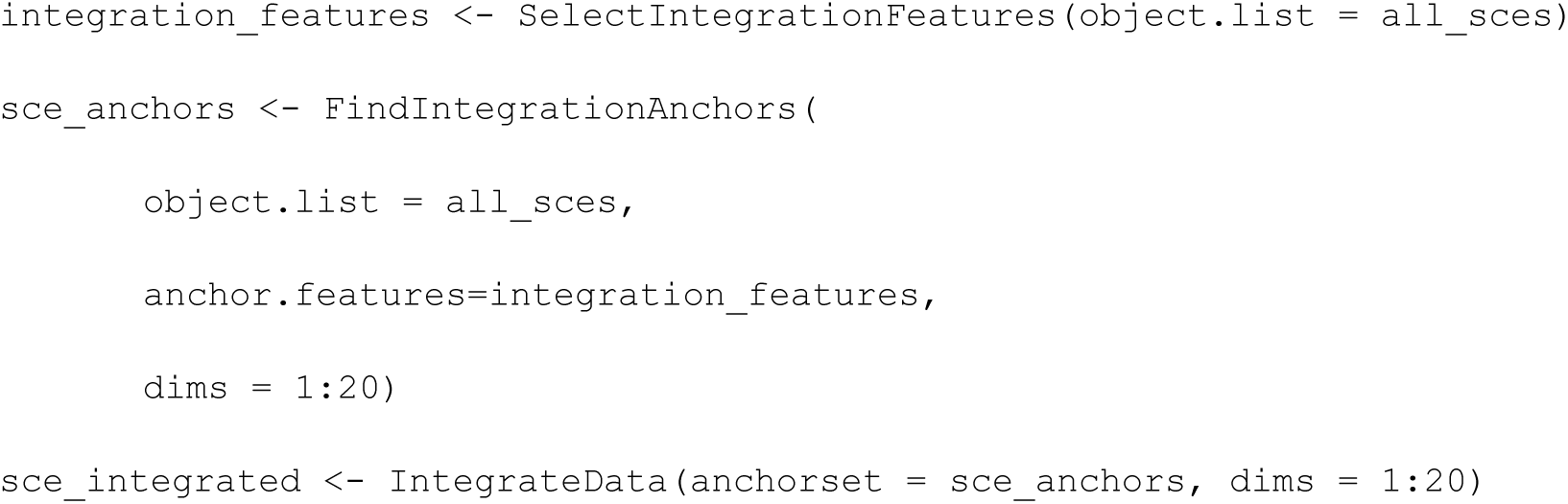

Number of PCA components considered on integrated data: 22

**5°) Analyzed data access:**

https://applisweb.vim.inrae.fr/Kodra

